# Genetic Diversity in Common Carp Stocks from a Natural Reservoir Assayed by Randomly Amplified Polymorphic DNA Markers

**DOI:** 10.1101/336669

**Authors:** Shahid Mahboob, K. A. Al-Ghanim, Norah M. A. Al-Mulhim

**Affiliations:** Department of Zoology, College of Science, King Saud University, P. O. Box 2455, Riyadh, Saudi Arabia

**Keywords:** Genetic diversity, common carp, freshwater fish, DNA markers

## Abstract

**Background:** The common carp *(Cyprinus carpio)* is a freshwater fish that is cultured throughout the world. *C. carpio* belongs to the family Cyprinidae, the largest family among freshwater teleosts, accounting for 10 % of total fish production. Specifically, the common carp is cultured in more than 100 countries in worldwide and accounts for over 3 million metric tons of total annual freshwater fish production. The population of common carp (C. *carpio*) is declining in freshwater reservoirs because of various human activities, such as overfishing, pollution, hybridization with domestic carp, and loss of breeding grounds because of habitat degradation.

**Results:** In total, 30 specimens were collected from each of four different locations (WH1, WH2, WH3 and WH4) in the reservoir. Five random decamer primers were used to assay polymorphisms within and between the population of *C. carpio.* A total of 60 bands were produced by these primers, out of which 50 bands were polymorphic and 10 bands were monomorphic. The mean highest polymorphism (100 %) was observed in the specimen collected from WHS4 stock, followed by 89.05, 87.62 and 76.66 % of the fish collection from WH3, WH2, and WH4, respectively. Nei’s genetic distance values ranged from 0.0006 to 0.1005. Highest and lowest genetic distance were 0.1005 and 0.006 in fish collected from WH1 and WH2, respectively. Average value of heterozygosity ranged from 0.3008 to 0.3748. A *C. carpio* UPGMA dendrogram was constructed to observe fish phylogeny. Phylogenetic clusters by RAPD indicated that fish stock of WH2, WH3, and WH4 were closely related to each other.

**Conclusion:** It was concluded that RAPD analysis can be successfully used as a marker to generate information regarding the percent homology within stock of common carp, which may be used to trace the progeny to the parents and is helpful for the improvement of breeding programs.

## 1. Introduction

The common carp *(Cyprinus carpio)* is a freshwater fish that is cultured throughout the world [1]. *C. carpio* belongs to the family Cyprinidae, the largest family among freshwater teleosts [2], accounting for 10 % of total fish production [3-4]. Specifically, the common carp is cultured in more than 100 countries in worldwide and accounts for over 3 million metric tons of total annual freshwater fish production [3]. This fish is omnivorous and feeds on phytoplankton, zooplankton, protozoans, and insect larvae to meet its requirements for growth [5]. The population of common carp (C. *carpio*) is declining in freshwater reservoirs because of various human activities, such as overfishing, pollution, hybridization with domestic carp, and loss of breeding grounds because of habitat degradation [6].

The genome of *C. carpio* was sequenced by combining data from various next-generation sequencing platforms [3]. Genetic diversity is important for wild stock and cultured fish species for the maintenance of evolutionary potential and the fitness of their population [3, 7]. It enhances growth potential, competitive potential, resistance against diseases, and also protects them against the threat of extinction. The decline in genetic variation in freshwater fish species may affect fitness and reduce its ability to adjust to the ever-changing environment [9-9]. The mutation, migration, and gene flow produce genetic variation in freshwater fish species and resultantly cause evolutionary changes in the fish [10]. Genetic diversity promotes the species′ capability to adapt to environmental changes and is important for the survival of species. Population genetic data have a wide range of applications in the field of management and conservation of natural resources, and that of genetic enhancement programs [11-12]. The conservation of allelic variation is important for retaining the genetic integrity and evolutionary potential in a natural population and allow adaptation to environmental changes [13]. The regular monitoring of riverine fisheries stock is required to avoid their genetic decline [14-15]. Before achieving fisheries restocking programs, it is necessary to examine the genetic diversity levels of the natural population [16]. The information associated with the genetic structure of wild fish may be helpful in restocking programs to monitor changes in genotype frequencies [17]. Aquaculture geneticists are now utilizing all the accessible modern technologies, such as DNA markers, gene sequencing, and genome mapping to interpret the mechanisms that lead to genetic variations in aquaculture [18]. The molecular markers are an important tool that has changed the genetic management of aquaculture [19]. DNA markers are the best current practice for population conservation at a sustainable level and for the construction of genetic linkage maps [11].

The randomly amplified polymorphic DNA (RAPD) technique is used for the investigation of genetic parameters, both in wild and inland fish stocks [20]. “Molecular markers such as Microsatellites, RAPDs, amplified fragment length polymorphism (AFLPs), restriction fragment length polymorphism (RFLPs) help the estimation of genetic diversity and are in practice for the conservation of fish population, parentage analysis, sex determination, identification of disease carriers, and transgenesis [21]”. In RAPD analysis, no prior information about the genetic content of the fish species is known, and this technique is currently used because it is fast and exposes a high degree of polymorphism [22]. This marker is also used to assess the effects of anthropogenic activities on genetic material [23-24]. Although common carp is commercially an important fish and its genetic data is relatively scarce [25]. The goals of this study were to document the genetic diversity and polymorphism in the common carp population of Riyadh River, Saudi Arabia through RAPD analysis, and also to highlight possible measures of conservation of such fish resources in a riverine template. In fact, the present investigation is the first report on the structure of the genetic diversity of *Cyprinus carpio* from Saudi Arabia.

## 2. Materials and Methods

### 2.1 Study Area

Wadi Hanefah is also known as Riyadh River/Riyadh Lake and has a length of 120 km (75 mi) from northwest to southeast, cutting through the city of Riyadh, Saudi Arabia. Riyadh City has a population of approximately 4 million. This water reservoir receives water treated by the Riyadh municipality’s sewerage system and untreated discharge from local industry and adjacent areas along the length of the river. The water of this reservoir is used for irrigation of various fruit farms and vegetables grown in adjacent areas [26].

### 2.2 Sampling of Fish

A total of 120 specimens of *Cyprinus carpio* was procured from four pre-determined sampling sites in the Wadi Hanefah (Riyadh River), Riyadh Saudi Arabia. The fish specimens were collected along the length of River Riyadh from Wadi Labn, Wadi Ubayr, Wadi Liha, and Al-Hair, which were designated as WH1, WH2, WH3, and WH4 (Fig 1). The total length of the specimens ranged from 87.00 ± 2.34 to 114.44 ± 4.62 cm. A blood sample from each stock fish was obtained from a caudal fin clipping vein using heparinised syringe and transferred to fresh tube with phosphate buffered saline and blood was preserved in 95 % ethyl alcohol at −20 °C. The specimens were labelled properly and transported to the Fisheries Lab for DNA extraction.

**Fig. 1.**
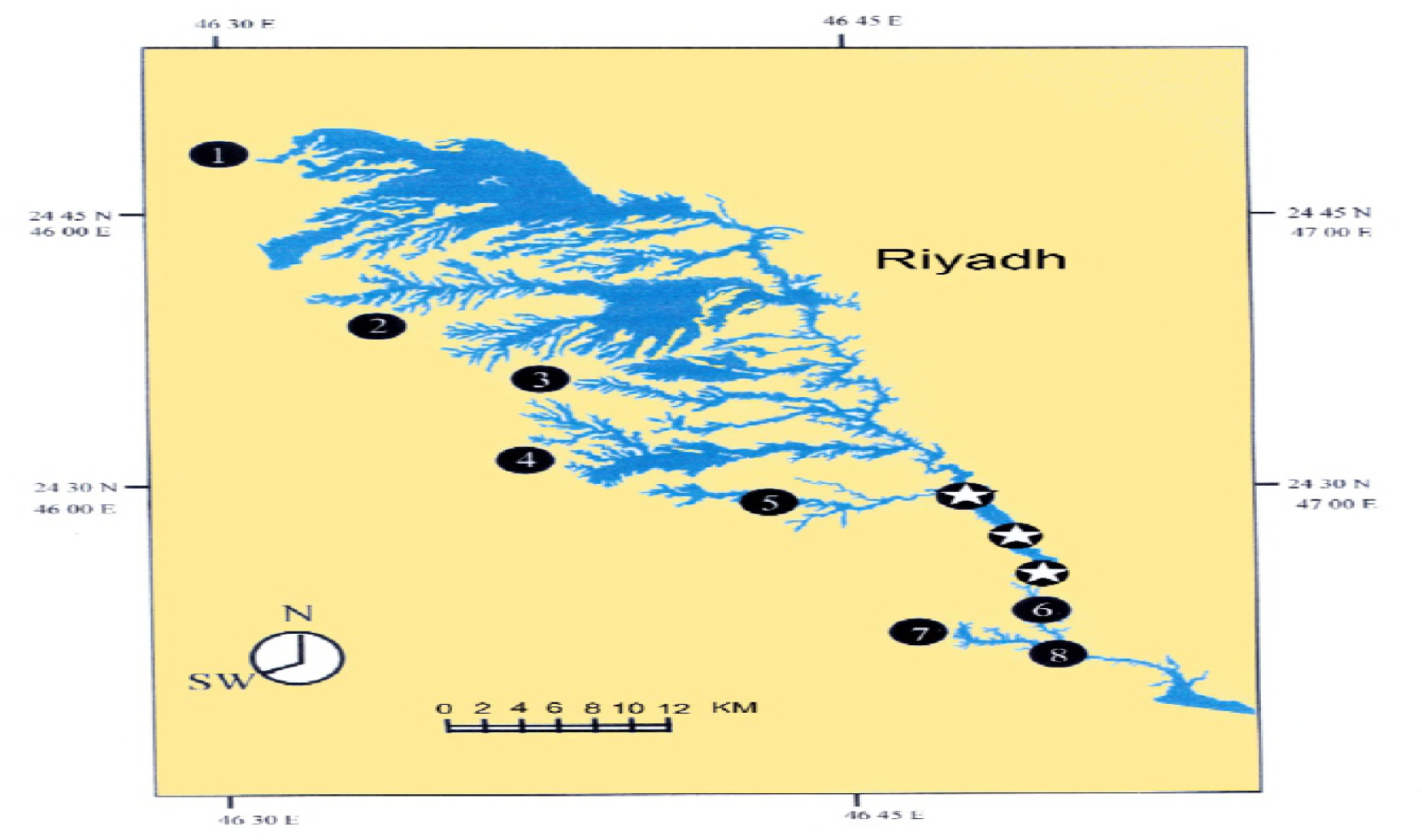
Map of Wadi Hanifah sampling stations. Site 1: Wadi Labn; Site 2: Wadi Ubayr; Site 3: Wadi Liha; Site 4: Al-Hair

### 2.3 Isolation of DNA

Genomic DNA was isolated by the procedure described by [27] with slight modification as described by [28]. DNA quality and quantity were assessed through agarose gels in a horizontal gel electrophoresis chamber utilizing spectrophotometric methods.

### 2.4 PCR Amplification of RAPD loci

Five random decamer primers (OPF-01, OPF-03, OPF-012, OPP-11, and OPP-16) were utilized in the detection of polymorphisms, which were attained from Operon Technologies. Extracted genomic DNA was amplified by polymerase chain reaction. Random decamer primers with their primer sequence and annealing temperature are presented in Table 1. The sequences of the primers were taken from the literature [29] and oligonucleotides were custom synthesized by Eurofins genomics, Canada. Specific parental band profiles were generated by these five primers in at least three replicate PCRs. For non-denatured gel electrophoresis, 40 % acrylamide gel was used. A mixture of 95 g acrylamide and 5 g N, N′ methane-bis-acrylamide was dissolved in double distilled H2O to a total volume of 250 mL at 37 °C. Five microliters of PCR products together with 1 µL loading dye, which was 0.5 blue dextran deionized bromide, and 3 µL DNA ladder were electrophoresed at a consistent voltage (280 v for 90 min). When the electrophoretic resolution of the PCR product was completed, the clips were gently removed and the gel was washed in silver staining solution.

**Table 1:** Random decamer primers with their Primer sequence, GC content, and annealing temperature

| Sr. No | Locus | Primer sequence(5'-3') | G + C (%) | Tm (C) |
| --- | --- | --- | --- | --- |
| 1 | OPF-01 | ACGGATCCTG | 60 | 32 °C |
| 2 | OPF-03 | CCTGATCACC | 60 | 32 °C |
| 3 | OPF-12 | ACGGTACCAG | 60 | 32 °C |
| 4 | OPP-11 | AACGCGTCGG | 60 | 34 °C |
| 5 | OPP-16 | CCAAGCTGCC | 60 | 34 °C |

### 2.5 Similarity Analysis

RAPD-ISSR data was used to generate a similarity matrix using the method of [30].

### 2.6 Data analysis

The genotypic data obtained from band counting were analyzed using the programs POPGENE and TFPGA. The genotype data for each locus were subjected to accurate analysis to estimate genetic diversity in the stock of *C. carpio.* The banding patterns generated by RAPD markers were scored on the basis of presence or absence of visible, clear, and reproducible bands. The presence of the band was scored 1 and the absence of the band was scored 0. The RAPD loci were utilized for determination of genetic diversity, the number of polymorphic loci, and genetic distance. They were also used to construct an unweighted pair group method for the arithmetic mean for the UPGMA dendrogram for the population using Nei’s unbiased distance [29].

For each sample, the proportion of polymorphic loci (P %), as well as the meaning of genetic diversity (H %) was calculated by POPGENE v.1.31. POPGENE software was utilized for analysis of polymorphic loci, genetic diversity within the population, genetic diversity between populations, and construction of a dendrogram based on Nei’s unbiased genetic distances. Genetic distances were estimated by utilizing TFPGA (Tool for population genetic analysis).

#### Ethical Guidelines

All the ethical guidelines defined by Ethical Committee of College Science, King Saud University animals research were followed. Clove oil sedative was used to euthanase the fish.

## 3. Results

### 3.1 RAPD analysis

A total of 60 bands was observed. Out of which, 50 bands were polymorphic and 10 bands were monomorphic (Table 2). The prevalence of bands differed greatly among the fish samples from different locations and use of primers. The maximum number of bands (14) was found in the fish collected from WH4 produced by primer OPF-03 and the lowest number of bands (9) in *C. carpio* was collected from WH1 by primer OPF-01.

**Table 2:** Polymorphic and Monomorphic bands

| Locus | No. of polymorphic bands | No. of monomorphic bands |
| --- | --- | --- |
| OPF-01 | 9 | 3 |
| OPF-03 | 15 | 4 |
| OPF-12 | 13 | 0 |
| OPP-11 | 13 | 0 |
| OPP-16 | 13 | 3 |
| Total | 50 | 10 |

### 3.2 Polymorphism among the Primers

The percentage of polymorphism in the *C. carpio* populations was as follows: OPF-12 (100.00 %) > OPP-11 (87.50 %) > OPF-01 (84.22 %) > OPF-03 (82.43 %) > OPP-16 (62.50 %). Five primers were used, and the highest and lowest polymorphism was observed at 100 and 64.85 % through OPF-12 and OPP-16 primers, respectively (Table 3). The highest (100 %) polymorphism by all the five primers was recorded in *C. carpio* harvested from WH3, followed by 88.55 %, 79.88 %, and 72.22 % from WH4, Wh2, and WHI, respectively (Table 3.).

**Table 3:**
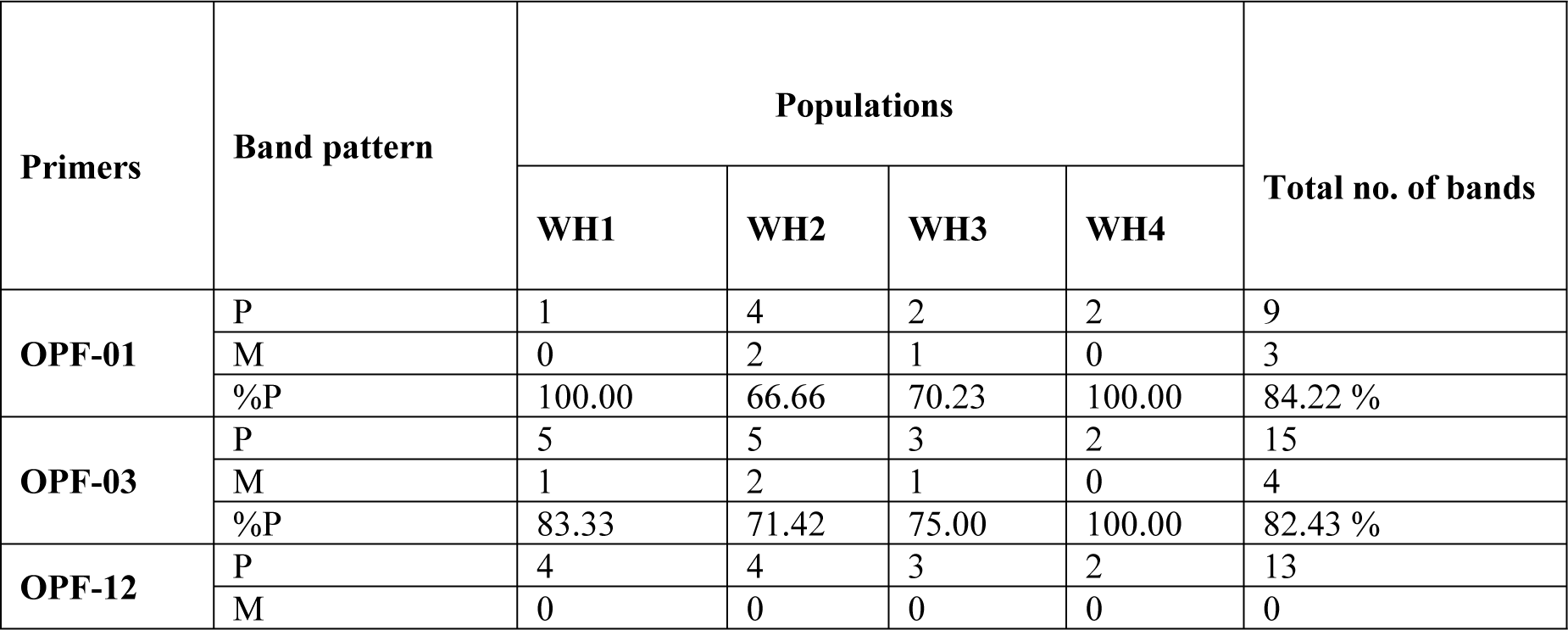
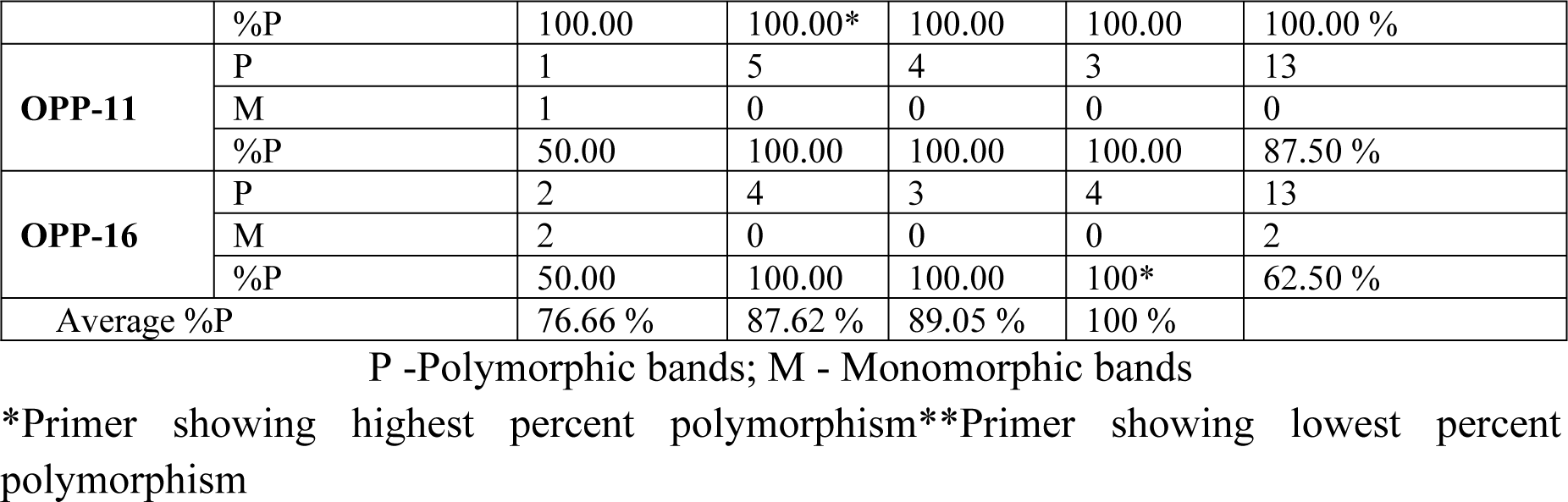
Total number of amplified fragments, number of polymorphic bands, and percentage polymorphisms generated by PCR using five primers

### 3.3 Genetic Diversity

The genetic diversity in the populations of *C. carpio* collected from the Riyadh River was estimated in terms of the number of alleles, heterozygosity, observed heterozygosity, expected heterozygosity, and average heterozygosity over five RAPD loci. The natural populations of *C. carpio* exhibited a decline in the genetic diversity because of anthropogenic activities, such as overfishing, pollution, deforestation, construction of dams, unintended selection of broodstock, genetic drift, and inbreeding effects.

### 3.4 Allelic frequency

Allelic frequencies for each locus in the four stocks of *C. carpio* is presented in Table 4. Null alleles were found in all populations of this fish collected from the four different locations. The value of a null allele was 1, which was found in all stocks in this river.

**Table 4:**
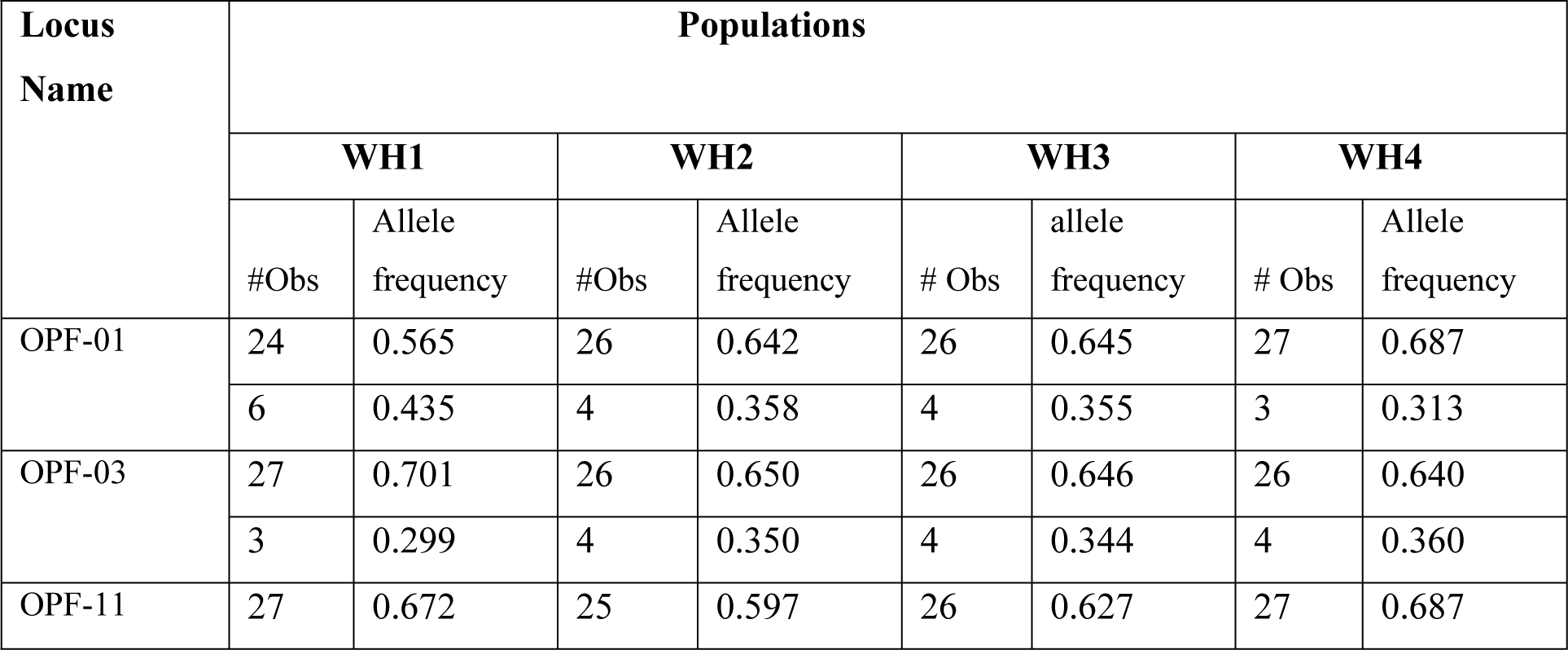
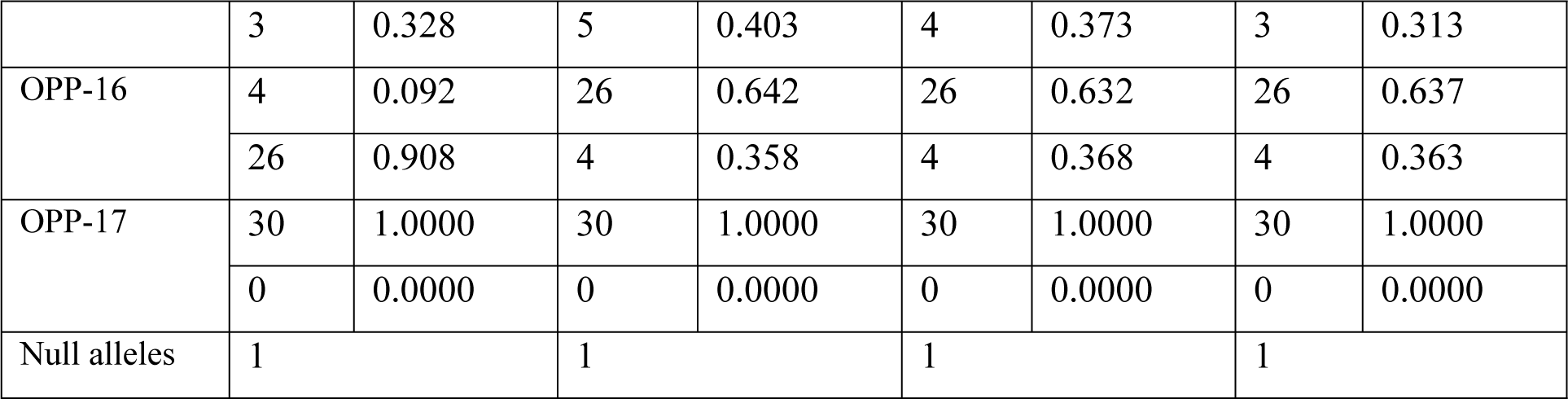
Allele frequencies in four populations of *C. carpio* at five loci

### 3.5 Heterozygosity

A low to moderate level of heterozygosity was estimated in the C. *carpio* population of this reservoir. The average value of heterozygosity ranged from 0.378 to 0.473. The heterozygosity was 0.378, 0.467, 0.462, and 0.473 in *C. carpio* collected from WH1, WH2, WH3, and WH4, respectively (Table 5). The highest level of heterozygosity was found in the fish collected from WH4. From all five loci, no negative values of heterozygosity were observed.

**Table 5:** Heterozygosity values in four populations of *Cyprinus carpio* at five loci hets: heterozygosity; het freq: heterozygosis frequency

| Locus name | Populations |  |  |  |  |  |  |  |
| --- | --- | --- | --- | --- | --- | --- | --- | --- |
|  | WH1 |  | WH2 |  | WH3 |  | WH4 |  |
|  | #hets | het freq | # hets | het freq | # hets | het freq | # hets | het freq |
| OPF-01 | 15.31 | 0.495 | 13.96 | 0.465 | 12.95 | 0.436 | 13.94 | 0.467 |
|  | 15.31 | 0.495 | 13.96 | 0.465 | 12.95 | 0.436 | 13.94 | 0.467 |
| OPF-03 | 13.33 | 0.431 | 13.97 | 0.467 | 13.93 | 0.466 | 13.93 | 0.469 |
|  | 13.33 | 0.431 | 13.97 | 0.467 | 13.93 | 0.466 | 13.93 | 0.469 |
| OPF-12 | 13.30 | 0.430 | 13.98 | 0.469 | 12.99 | 0.437 | 14.54 | 0.487 |
|  | 13.30 | 0.430 | 13.98 | 0.469 | 12.99 | 0.437 | 14.54 | 0.487 |
| OPP-11 | 4.83 | 0.158 | 13.97 | 0.468 | 13.94 | 0.469 | 13.95 | 0.470 |
|  | 4.83 | .158 | 13.97 | 0.468 | 13.94 | 0.469 | 13.95 | 0.470 |
| OPP-16 | 0.000 | .000 | 0.000 | 0.000 | 0.000 | 0.000 | 0.000 | 0.000 |
|  | 0.000 | .000 | 0.000 | 0.000 | 0.000 | 0.000 | 0.000 | 0.0000 |
| Average Heterozygosity | 0.378 |  | 0.467 |  | 0.452 |  | 0.473 |  |

### 3.6 UPGMA Cluster Analysis

Major clusters generated by the UPGMA dendrogram are presented in Figure 4.3. The phylogenetic tree showed the construction of three clusters. The first cluster was formed between the WH4 and WH3 populations, the second cluster was between the WH2 and WH3 populations, and the WH1 population formed a separate cluster alone. Genetic distance was highest among the populations of WH1and WH3, which indicated a heterozygous genotype, whereas the lowest was estimated between the HM and HK populations, which indicated a homogenate genotype. Node distance included population (1) 0.0006, 2, 3 (2) 0.0024 2, 3, 4, and (3) 0.0995, 1, 2, 3, 4.

**Fig.2:**
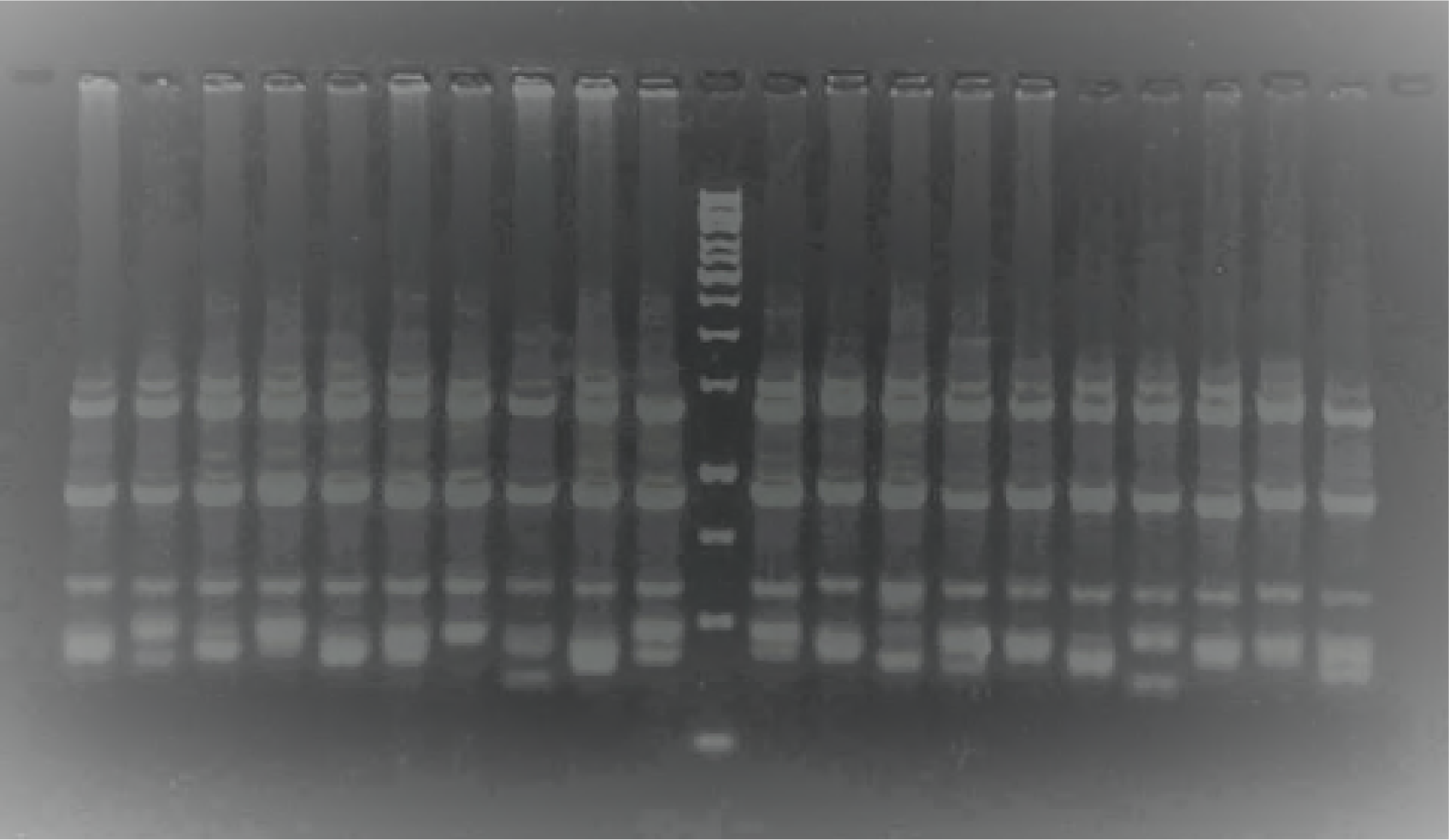
RAPD gel profile of *Cyprinus carpio* (first 10 lanes) from WH1 and (second 10 lanes) from WH2. Marker: 1 Kb marker with selected primers.

**Fig. 3.**
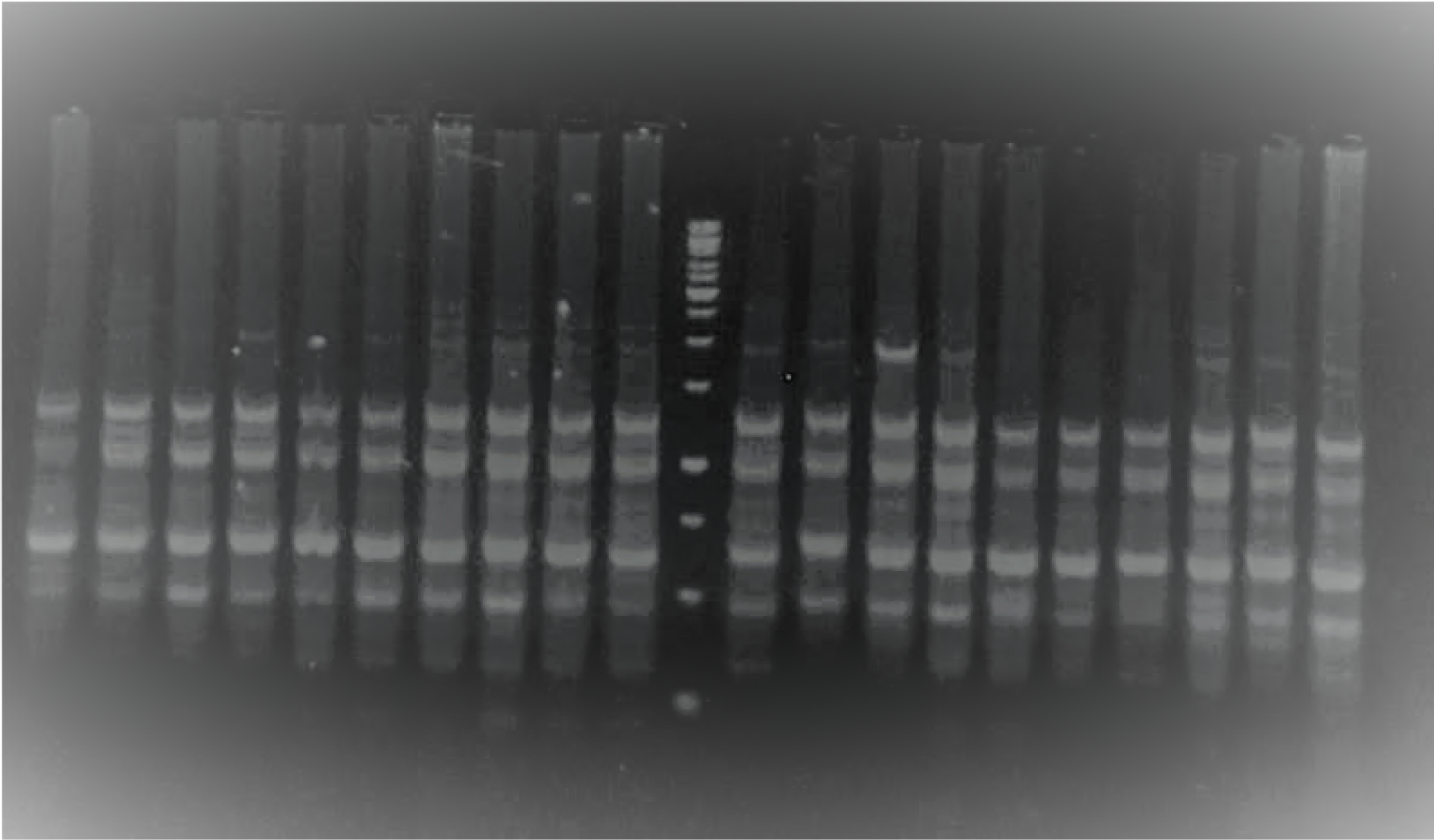
RAPD gel profile of *Cyprinus carpio* (first 10 lane) from WH3 and (second 10 lane) from WH4. Marker: 1 Kb marker with selected primers.

**Fig. 4:**
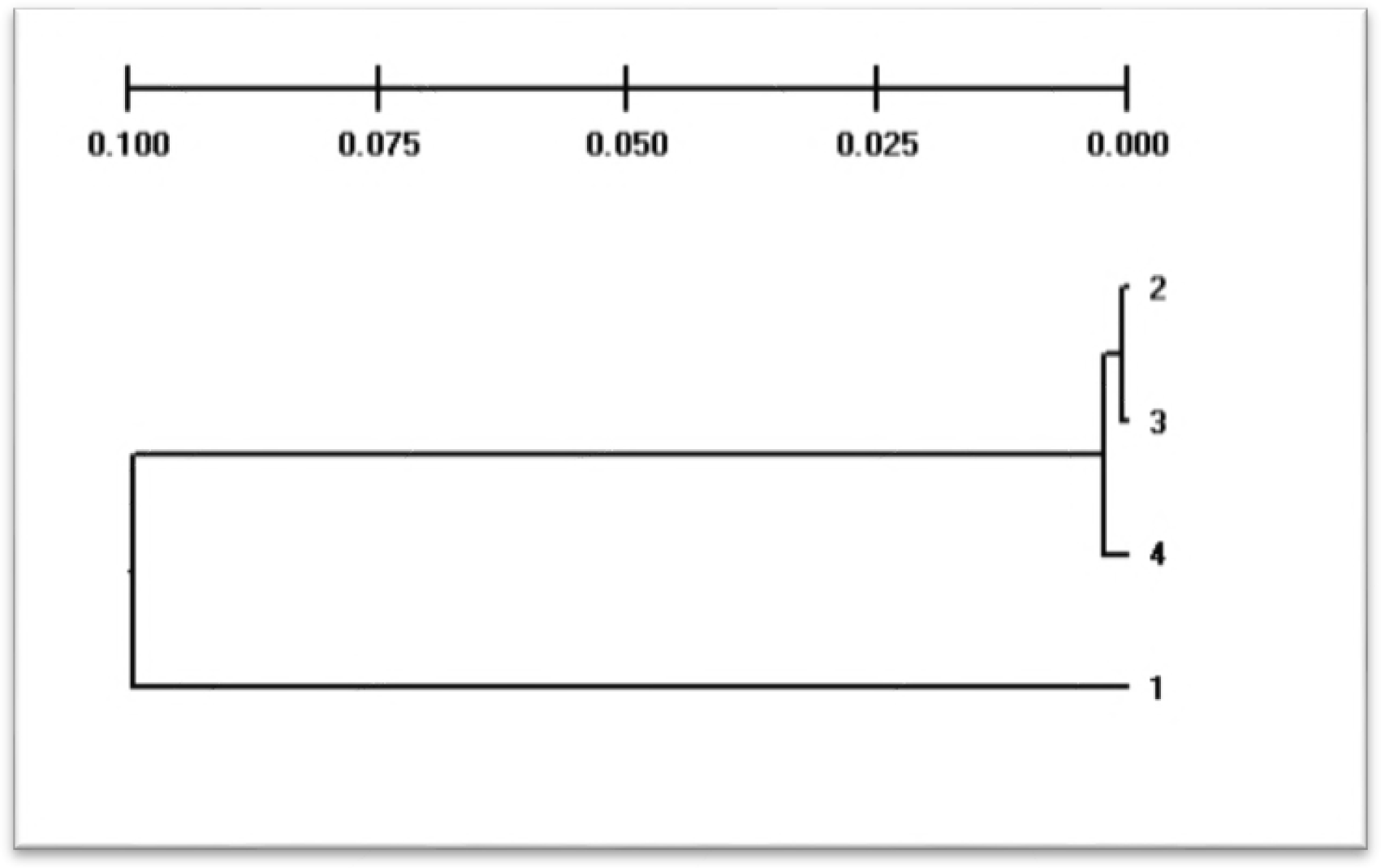
Dendrogram construction based on Nei’s genetic distance among *Cyprinus carpio* populations

## 4. Discussion

*C. carpio* collected from WH1, WH2, and WH3 exhibited the highest level of genetic diversity (1.0000) produced by the primers OPF-01, OPF-12, and OPP-11. The species introgression was caused by anthropogenic activities and seasonal flooding for this large genetic diversity. Although *C. carpio* is one of the most highly consumed fish worldwide, information regarding the genetic diversity of farmed and wild stocks is still limited in the literature. Isoenzyme analysis of *C. carpio* samples exhibited genetic variation between stocks originating from the same geographic area of this water reservoir [31]. The highest and lowest polymorphism were observed at 100 and 64.85 % through OPF-12 and OPP-16 primers, respectively (Table 2). These percentages were higher than those reported by [32]. They observed an 86.00–92.11 % polymorphic loci ratio in five species of snappers using the RAPD technique. In this study, polymorphisms of alleles were comparatively higher in comparison with findings of other studies. Five random decamer primers produced both monomorphic and polymorphic bands. A total of 60 bands were produced and of these, 50 bands exhibited polymorphism. [22] assessed the genetic variation of six stocks of *C. carpio*, which were collected from India, Indonesia, and Vietnam by using RAPD markers. Eight decamer primers were chosen for estimation of common carp genotypes, which included OPA-7, OPA-20, OPB-17, OPF-10, OPF-9, OPG-4, OPG-9, and OPP-16 primers. A total of 492 bands was recorded, out of which 57.1 % were polymorphic. [22] analyzed the genetic variation of *Labeo fimbriatus* by using eight random primers, which included OPA-07, OPP- 11, OPB-17, OPF-01, OPF-03, OPF-12, OPD-19, and OPP-16 and all primers produced polymorphic bands. In this study, higher genetic diversity was recorded within the *C. carpio* harvested from WH4 (1.000), and lower genetic diversity (76.66 %) from WH1. This means that common carp from WH4 had a higher proportion of heterozygous genotypes compared to that of the other three locations. The results of this research work were consistent with the results of [33].

Genetic distance ranged between 0.0006 to 0.1005. The highest and lowest genetic distances were 0.1005, and 0.006 in the fish collection from WH1 and WH2, respectively. [34] examined the genetic distance and Shannon′s information index values and provided evidence of differences in genetic diversity among five populations of *Megalobrama amblycephala*. Heterozygous genotypes were also estimated by RAPD markers in *C. carpio* populations and the highest proportion of heterozygous genotypes was exhibited in the fish from WH1. [35] mentioned that genetic distance values above 0.3 differentiated most fish species. The results of this research indicated a low level of genetic variation (0.18) among the stocks of common carp collected from different locations in the Riyadh River [36]. However, our findings were not in line with the results of [31], who reported variation ranging from 0.256 to 0.112. Three clusters were constructed and the WH3 population was closely related to WH4 population, the WH2 population was closely related to WH3 stock, and WH1 was genetically distant from all other stocks. [22] reported a grouping in three different populations of *Labeo fimbriatus* into two clusters. [25] analyzed whole brood stocks of two Hungarian common carp farms and no groupings were observed in the dendrogram. [37] assessed the genetic similarity in common carp genera and its phenotypes, used 13 random decamer primers, and observed a closer proximity to bighead carp. [38] analyzed the genotype of *Catla, Labeo rohita*, and *Cirrhinus mrigala* by using RAPDs and a highly reproducible RAPD profile was detected among the fish species. We are of the opinion that the anthropogenic activities and the discharge of untreated waste from the adjacent areas of Riyadh River are a big challenge to the diversity of common carp. Immediate attention is required to conserve its populations and their diversity. The loss of genetic diversity in fish may reduce the fish’s ability to cope with environmental changes [39].

## Conflict of Interest

Noting to declare.

## Acknowledgements

The authors (SM and KAAG) would like to express their sincere appreciation to the Deanship of Scientific Research at the King Saud University for its funding of this research through the Research Group Project No. RG-1435-012. The authors also thank the Deanship of Scientific Research and RSSU at King Saud University for their technical support.

